# Roles of Polycomb gene *EED* in pathogenesis and prognosis of acute myeloid leukemia and diffuse large B cell lymphoma

**DOI:** 10.1101/444745

**Authors:** Wenhua Yu, Haiwei Du

## Abstract

In this study, we performed correlation analysis of polycomb gene *EED* and hematologic malignancies using the omics and clinical data of acute myeloid leukemia (LAML) and diffuse large B-Cell lymphoma (DLBC) from TCGA database. We found that: (1) High *EED* mRNA level was associated with poor prognosis and high CALGB cytogenetics risk of LAML patients. (2) *EED* mRNA level in DLBC cancer cells was higher than control cells. (3) *EED* gene expression could be regulated by both copy number alterations and DNA methylation. (4) Additionally, there were different *EED* co-expression genes nets in the two kinds of hematologic malignancies. In all, we confirmed that there are potential clinical significance of *EED* gene in pathogenesis and prognosis of hematologic malignancies.

## Introduction

Polycomb repressive complex 1 and 2 (PRC1 and PRC2) are canonical chromatin silencing related protein complex created by different PcG gene assemblies, which can maintain the transcriptional repression of target genes through H3K27me3 and H2AK119Ub chromatin modification. Abnormal Expression and inactivation of Polycomb group (PcG) genes have been considered as important causes of hematological malignancies including acute myeloid leukemia (LAML) and diffuse larger B-cell lymphoma (DLBC) due to the derepression of pro-oncogenes (Iwama 2017). Embryonic Ectoderm Development (EED) protein is the core component of Polycomb group protein family, which is the essential PCR2 structural protein and has direct interaction with PRC1 (Cao et al. 2014; Margueron & Reinberg 2011). Besides, it cloud regulate gene expression by interaction with histone deacetylases (Ai et al. 2017). So, we guess that *EED* gene could play vital roles in hematopoiesis and hematological malignancies.

LAML is characterized by ectopic proliferation of myeloblasts in the bone marrow with highly clinical heterogeneity. LAML is defined as 8 subtypes (M0-M7) according to the French-American-British (FAB) classification (Bennett et al. 1976). LAML is frequently occurred in older adults, but only 10-20% LAML patients with more than 60-year old possess more than 5 year overall survival after receiving cytarabine-based chemotherapy or hematopoietic stem cell transplant treatment (Yang & Wang 2018). DLBC is the most prevalent non-Hodgkin lymphoma (NHL) in adulthood with high malignancy and aggressiveness. DLBC is genetically heterogeneous, the somatic mutation identity among different DLBC patients is only 10-20%.

The roles of PcG genes in LAML are elusive. High expression of *EZH2* gene, coding PRC2 catalytic subunit, is associated with extramedullary infiltration in LAML and *EZH2* silencing suppresses cell proliferation, cell migration and increases cell apoptosis in LAML through inhibiting *MMP2* expression and increasing *E-cadherin* expression (Zhu et al. 2016). However, lower *EZH2* expression also correlates with poorer prognosis of LAML patients due to loss of *EZH2* inhibits the expression of *HOX* gene and results in enhancing chemo-resistance of LAML cell lines (Gollner et al. 2017). Additionally, LAML patients with *EZH2* mutations predict unfavorable prognosis with worse overall survival and relapse-free survival (Saygin et al. 2018).

In DLBC cancer cells, the high H3K27me3 level is significantly correlated with EZH2 protein level and predicts inferior overall survival (Oh et al. 2014), and high level of H3K27me3 in bone marrow resident cells was also correlated with poor progression-free survival and overall survival of DLBC patients (Oh et al. 2016; Oh et al. 2014). However, significant correlation between single PRC2 gene and DLBC prognosis have not been reported up to now.

To our knowledge, the corresponding published articles reporting correlation among *EED* gene expression and prognosis of hematologic malignancies is absent. In our previous study, we found that conditional knockout of *Eed* gene impairs fetal hematopoiesis including erythropoiesis and formation of hematopoietic progenitor and stem cells (Yu et al. 2017). Accumulating evidence in mice model demonstrated loss-of-function mutations of *Eed* gene might be potential etiology of leukemia. Ikeda et al., report that *Eed* haploinsufficiency induces hematopoietic dysplasia and *Eed* heterozygous mice is susceptible to malignant transformation and developed leukemia in cooperation with *Evi1* over-expression(Ikeda et al. 2016). Ueda et al., demonstrate that *EED^I363M^* heterozygotes increases in the clonogenic capacity and bone marrow repopulating activity of hematopoietic stem/progenitor cells and is susceptible to leukemia through depressing *Lgals3* gene expression, encoding a multifunctional galactose-binding lectin (Ueda et al. 2016). Shi et al., report that suppression of PRC2 core genes *Eed*, *Suz12* or *Ezh1/Ezh2* results in proliferation arrest and differentiation of leukemia cells(Shi et al. 2013).

Based on our previous study and current articles, we propose the hypothesis that EED expression might correlate with the pathogenesis or prognosis of LAML and DLBC patients. In this study, omics data including transcriptome profiling, DNA methylation, single nucleotide polymorphism, copy number variant and clinical data of LAML and DLBC patients were retrieved from TCGA database, aiming to investigate prognostic significance of *EED* in LAML and DLBC patients, and preliminarily decipher the potential pathogenesis mechanism of *EED* involved in hematologic malignancies.

## Materials and Methods

### Patients and omics data

Total of 200 LAML patients and 48 DLBC patients were retrieved from The Cancer Genome Atlas (TCGA) data portal. The clinical data consisted of age, gender, history of neoadjuvant treatment, cancer and leukemia group B (CALGB) cytogenetics risk category, living status, clinical stage, bone marrow involvement, recurrence status, recurrence-free survival (RFS), overall survival (OS).

All the level 3 omics data including RNA-sequencing, single nucleotide polymorphism, copy number data, DNA methylation data and clinical data of LAML patients and DLBC patients used in this study were collected from TCGA database (https://cancergenome.nih.gov/) by UCSC Xena (https://xenabrowser.net/) and cBioPortal (http://www.cbioportal.org/) browsers (Cerami, et al 2012, Gao, et al 2013). Gene expression data of normal paired tissues used in this study were collected from the GTEx database (https://www.gtexportal.org/) by GEPIA browser (http://gepia.cancer-pku.cn/) (Tang, et al 2017). ChIP-seq data were collected from GEO database (https://www.ncbi.nlm.nih.gov/geo/).

The difference of mRNA expression level of *EED* between LAML and normal, between DLBC and normal was investigated. The difference of *EED* copy number and DNA methylation status among LAML patients was detected. The patients with different hematological malignancies were segregated into high *EED* mRNA level group and low *EED* mRNA level group by the cut-off values calculated by Youden index method from the receiver operating characteristic (ROC) curve (Bantis et al. 2014; Luo & Xiong 2013).

### Survival analysis

In LAML, 149 patients had both of survival information and *EED* expression data were incorporate into the survival analysis. In DLBC, 47 patients had both of OS information and *EED* expression data were incorporate into the OS survival analysis, and 45 DLBC patients had both of RFS information and *EED* expression data were incorporate into the RFS survival analysis. Some patients were excluded from survival analysis due to absent record of survival information or *EED* expression. The correlation analysis between *EED* expression level and OS or RFS times of patients with LAML or DLBC was calculated with log-rank test of Kaplan-Meier survival analysis.

### *EED* co-expressed genes screening

In our work, pearson and spearman scores among the expression of EED and other genes in respective LAML patients and DLBC patients was calculated, those genes with pearson and spearman scores > 0.4 or < -0.4 were identified as EED co-expressed genes. The KEGG signaling pathways of EED co-expressed genes were enriched, and the correlation network among EED co-expressed genes and signaling pathways were visualized by Cytoscape 3.6.1 software (http://www.cytoscape.org/).

### Statistical analysis

Gene expression differences among groups were analyzed by Mann Whitney test (two groups) or Kruskal-Wallis test (multiple groups) using GraphPad Prism 6 software. Univariate analysis and Multivariate Cox regression was used to analyze the risk factors of OS of LAML patients and DLBC patients. The difference of discrete data in groups was calculated by chi-square test or Fisher’s exact test. The continuous data was representing by mean ± standard deviation (SD). ^*^ indicated *p* < 0.05, ^**^ indicated *p* < 0.01, ^***^ indicated *p* < 0.0001.

## Results

### EED mRNA level were upregulated in DLBC patients

According to the TCGA and GTEx data combined analysis, we found that EED mRNA level were significantly upregulated more than 10 fold in DLBC cancer cells compared with the paired normal cells (*p* < 0.01, Figure 1A) and EED expression had the down-regulated tendency in LAML compared with normal samples, but there none of statistical difference. In contrast, conditional deletion of EED gene in hematopoietic cells could reduce the amount of B cells in E17.5 mouse fetal livers (Xie, et al 2014). It seems that EED is essential to the B cell proliferation and EED overexpression could be a risk factor of DLBC pathogenesis. Additionally, according to the UCSC Xena analysis, we found that EED mRNA level in DLBC cancer cells were significantly higher than LAML cancer cells (*p* < 0.0001, *t*-test, Figure 1B), it indicated that EED gene could have different expression patterns and functional significances in DLBC and LAML cancer cells.

**Figure 1.**
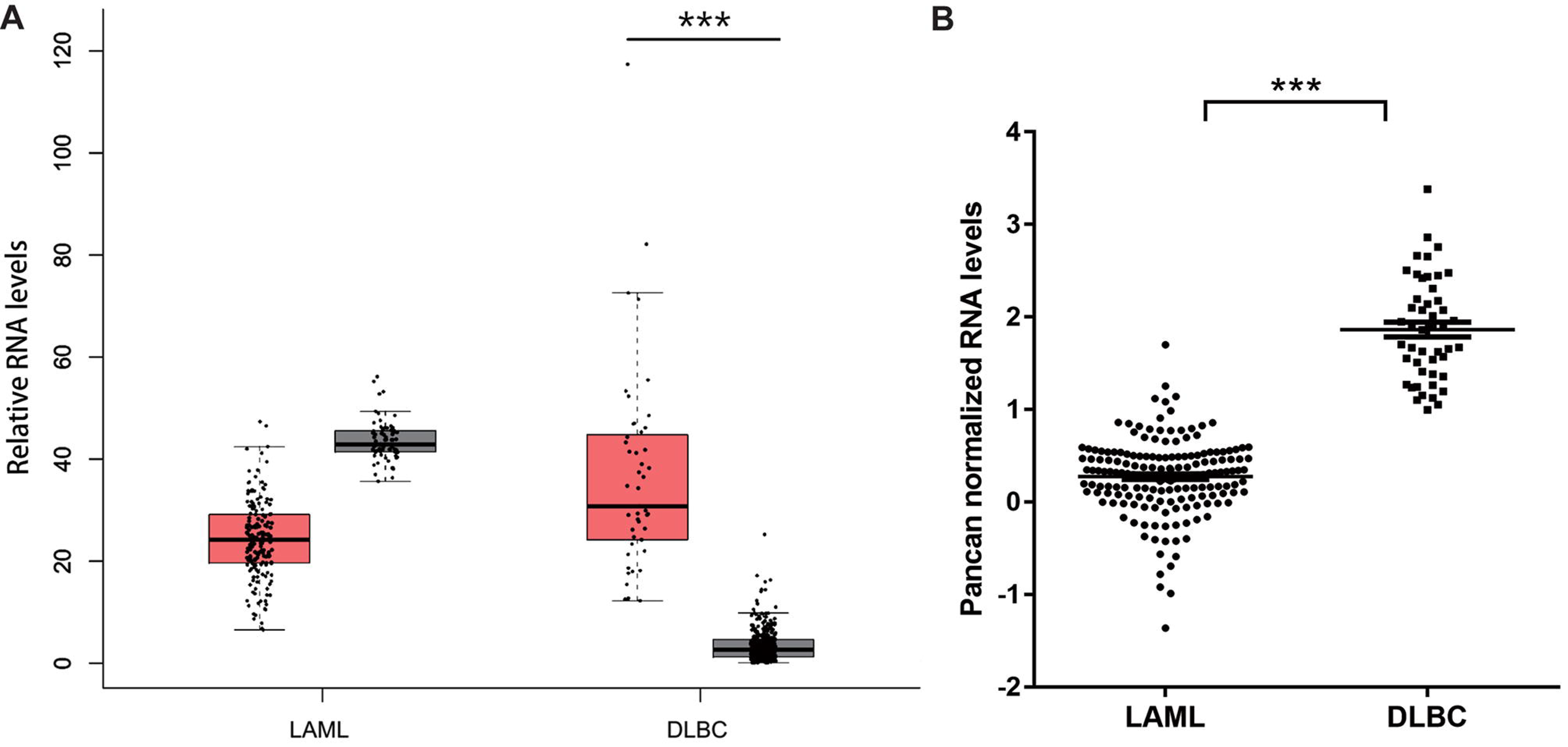
*EED* mRNA levels in cancer cells of LAML and DLBC patients. (A) Box plot of *EED* mRNA levels in hematologic malignancies and paired normal tissues was created by GEPIA Browser. Red and black box-plot respectively indicated hematologic malignancies and normal samples. (B) Dot plot of *EED* mRNA levels in cancer cells of patients with primary LAML and DLBC was created by GraphPad Prism 6 software. ^***^ indicated *p* < 0.001.

However, we did not find that the statistical association between EED mRNA level and the overall survival (OS), between EED level and recurrence-free survival (RFS) times of DLBC patients through Kaplan-Meier survival analysis (Figure S1B, S1C). And, we did not find the statistical association between EED mRNA level and the clinicopathological characteristics (such as age, gender, bone marrow involvement, clinical stage, and recurrence status) of DLBC patients (Table S1). Besides, the EED mRNA level was not statistical difference between patients with clinical stage I and II and patients with clinical stage III and IV. In the univariate analysis, we also did not find the statistical association between EED mRNA level and the OS, between EED level and RFS times of DLBC patients (Table S2).

### EED mRNA level were associated with LAML risk and prognosis

In spite of EED mRNA level did not changed obviously in LAML cancer cells (Figure 1A), we found that EED mRNA level in LAML-M3 subtype (Figure 2C, acute promyelocytic leukemia, APL) were significantly lower than other LAML subtypes (M0-M4, *p* = 0.0005, Kruskal-Wallis test). APL patients were more susceptible to hemorrhage than other LAML subtypes, which could be caused by vascular invasion of APL cancer cells containing lower EED mRNA level. It seems that lower EED mRNA level could increase the metastasis and invasion potential of APL cancer cells. Viewed together with this hypothesis, we found that haploinsufficiency of EED gene in mouse caused by conditional deletion or mutant knockin could also increase the metastasis and invasion risk of leukemia cells (Ikeda et al. 2016; Ueda et al. 2016).

**Figure 2.**
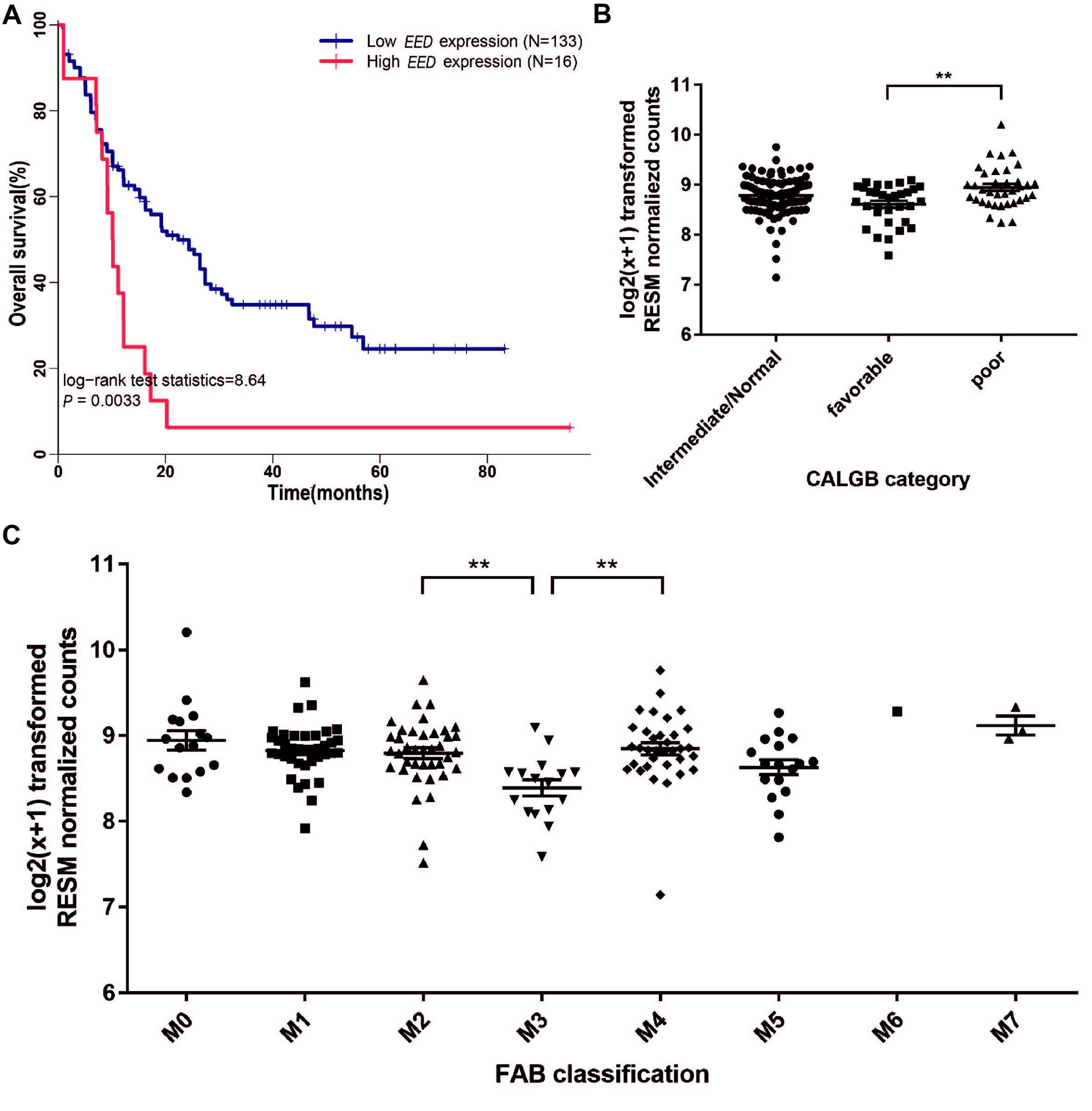
The association of *EED* mRNA levels and prognosis of LAML patients. (A) The association analysis of *EED* mRNA levels and overall survival (OS) time in LAML patients. (B-C) The association analysis of *EED* mRNA levels in different CALGB (B) and FAB (C) categories of LAML patients.

EED mRNA level were also associated with cancer and leukemia group B (CALGB) cytogenetics risk category of LAML, for EED mRNA level in the high risk (poor) group were significantly higher than the low risk (favorable) group (Figure 2B, *p* = 0.0038, Mann Whitney test). And, in the Kaplan-Meier survival analysis, we found that LAML patients with high EED mRNA level had worse prognosis than LAML patients with low EED mRNA level (Figure 2A, *p* = 0.0033, Log-rank test).

Besides, we found that there was significantly statistical association between EED mRNA level and some clinicopathological parameters (including CLAGB cytogenetics risk category and recurrence status) of LAML patients (*p* < 0.01, Table 1). And, in the univariate and multivariate analysis, we also found EED mRNA level, similar with age and CLAGB cytogenetics risk category, had significantly statistical association with the OS times of LAML patients (*p* < 0.01, Table 2). These results indicated that EED mRNA level, which similar with CALGB risk category, could be an independent LAML prognostic factor.

**Table 1.** The association analysis of EED mRNA level and the clinicopathological characteristics of AML patients.

| Parameters |  | EED mRNA level |  | P value |
| --- | --- | --- | --- | --- |
|  |  | High (N = 16) | Low (N = 133) |  |
| Age at initial pathologic diagnosis (Mean ± SD) | | 55.38 ± 18.02 | 54.59 ± 16.22 | 0.858 |
| Gender | Female | 3 | 66 | 0.031* |
|  | Male | 13 | 67 |  |
| History of neoadjuvant treatment | With | 2 | 34 | 0.359* |
|  | Without | 14 | 99 |  |
| Living Status | Living | 1 | 56 | 0.005* |
|  | Dead | 15 | 77 |  |

Note: \* indicates Fisher's exact test used in the significance analysis.

**Table 2.** Univariate and multivariate analysis of overall survival (OS) in AML patients.

| Parameters | Univariate analysis |  |  |  | Multivariate analysis |  |  |  |
| --- | --- | --- | --- | --- | --- | --- | --- | --- |
|  | P value | HR | 95%CI (lower/upper) |  | P value | HR | 95%CI (lower/upper) |  |
| Age at initial pathologic diagnosis (> 55 vs. ≤ 55) | <0.001 | 2.809 | 1.794 | 4.400 | <0.001 | 2.895 | 1.842 | 4.552 |
| Female vs. Male | 0.817 | 1.050 | 0.696 | 1.583 | 0.297 | 1.261 | 0.816 | 1.949 |
| History of neoadjuvant treatment | 0.044 | 1.605 | 1.012 | 2.545 | 0.016 | 1.766 | 1.111 | 2.808 |
| EED mRNA level High vs. Low | 0.004 | 2.269 | 1.293 | 3.984 | 0.002 | 2.590 | 1.419 | 4.729 |

Notes: HR indicates hazard ratio; CI indicates confidence interval.

### Genetic and epigenetic factors involved in the EED mRNA level changes

To analyze the causes of EED mRNA level changes, we searched the genetic variation of EED gene in our study population at first. Unfortunately, only one missense mutation (p.Leu196Gln) was found in the LAML group, which is located on the WD40 domain of EED protein and could impair the binding of EED protein and EZH2 protein. However, copy number alterations of EED gene had been found both in DLBC and LAML patients, and the subjects with increased copy number alterations of EED gene showed significantly higher EED mRNA level than the subjects with normal diploid of EED gene (Figure 3A and 3B). However, there was no directly statistical association between EED copy number additions and survival times of LAML and DLBC patients (Figure S2), EED mRNA level could be regulated both in genetics and epigenetics factors beyond itself alterations of EED gene.

**Figure 3.**
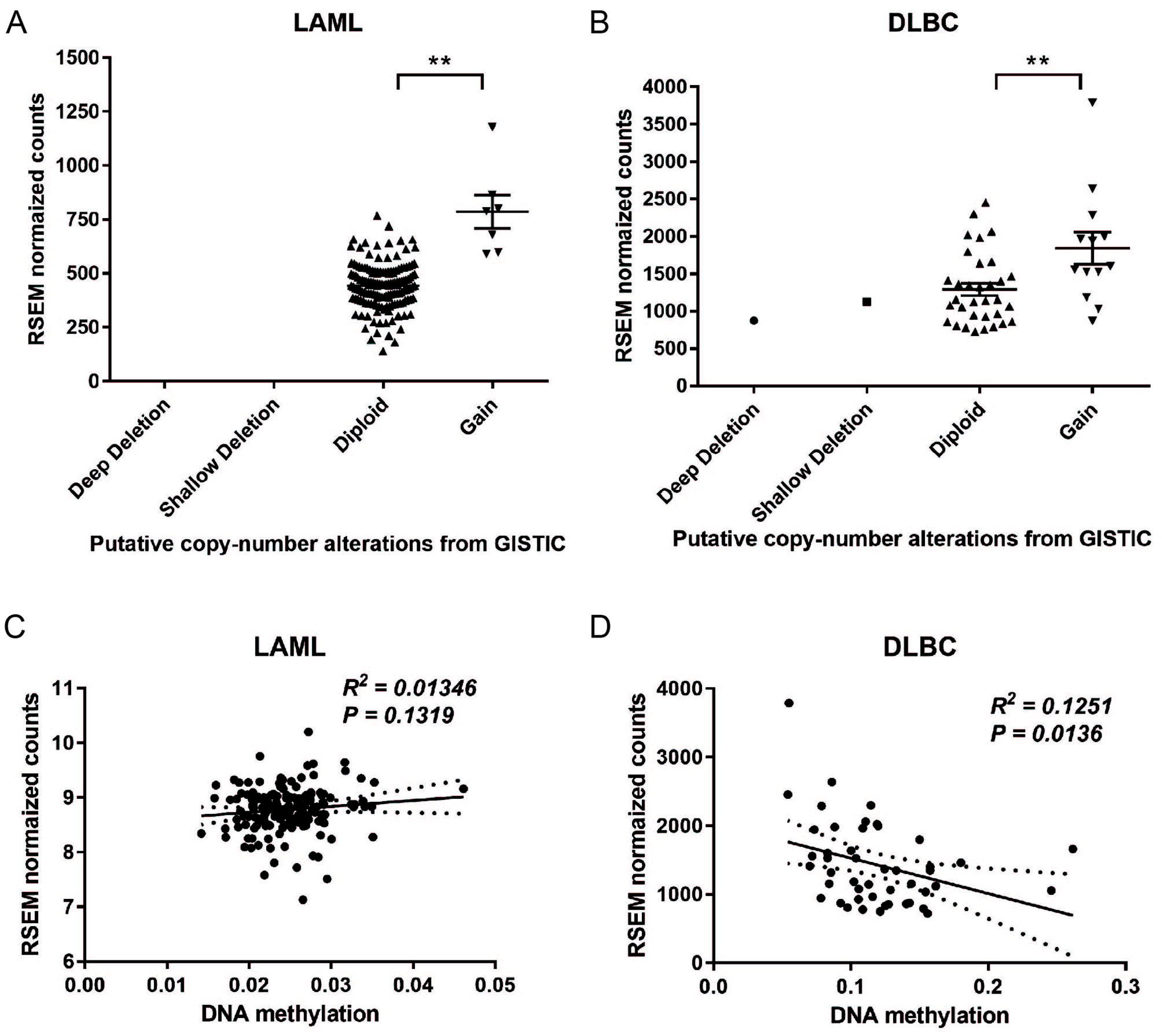
Association analysis of copy number alterations, DNA methylation and mRNA level changes of *EED* gene in LAML and DLBC cancer cells. (A-B) The association analysis of *EED* mRNA levels and copy number alterations in LAML (A) and DLBC (B). (C-D) The linear regression analysis of *EED* mRNA expression and DNA methylation in LAML (C) and DLBC (D) patients.

In the epigenetics analysis, we found that the DNA methylation status of EED gene were obviously different between LAML and DLBC cancer cells (Figure S3), and DNA methylation level of EED gene was positive associated with EED mRNA level in DLBC instead of LAML cancer cells (Figure 3C-D).

### The *EED* related regulatory networks in DLBC and LAML patients

To study the differences of EED mediated gene regulatory networks in LAML and DLBC cancer cells, we Filtered out the EED co-expression genes in LAML and DLBC cancer cells respectively by person and spearman scores (both absolute values was not lower than 0.4). Accounting to the results, there were 1069 EED co-expression genes in LAML cancer cells, but only 54 genes were co-expressed with EED gene in DLBC cancer cells, and there were only 2 co-expression genes (XRCC5 and ZNF252P) were shared among LAML and DLBC groups (Figure 4A). Those results indicated that there could are different EED related gene regulatory networks in DLBC and LAML cancer cells. Besides, it was interesting that there were only 80 EED co-expression genes were negative correlation with EED genes in LAML cancer cells.

**Figure 4.**
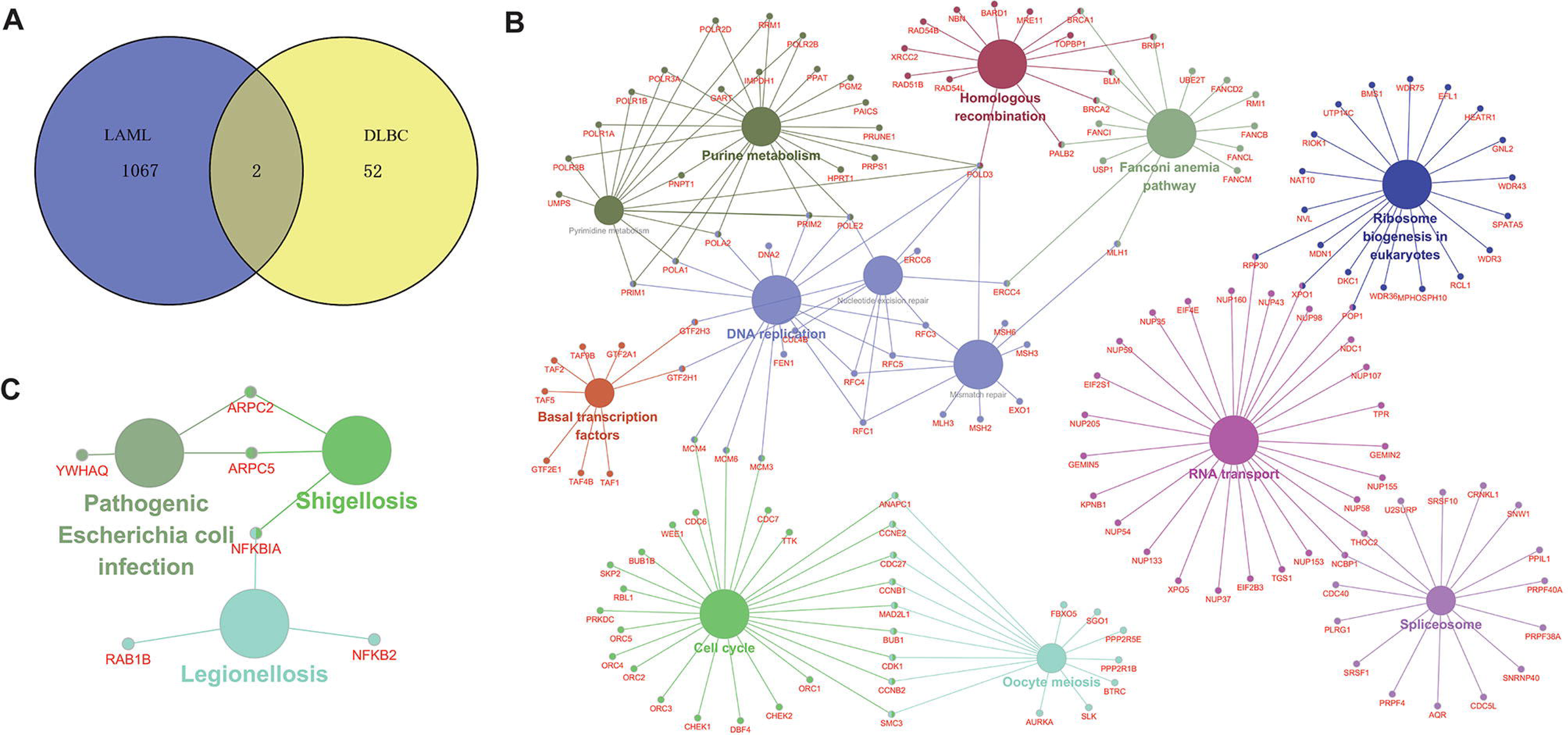
KEGG pathway analysis of the *EED* co-expressed genes in LAML and DLBC cancer cells. (A) The overlapping of *EED* co-expressed genes in LAML and DLBC cancer cells. (B-C) KEGG pathway analysis of *EED* co-expressed genes in LAML (B) and DLBC (C) cancer cells.

KEGG pathway analysis of the EED co-expressed genes showed that, EED gene could regulated the LAML pathological processes mainly through the cell proliferation and chromatin homeostasis related pathways, such as cell cycle, DNA replication, Fanconi anemia, homologous recombination, nucleotide excision repair, and mismatch repair (Figure 4B and Table S3); but in DLBC cancer cells, EED could regulated the immunological response to the bacterial infections (Figure 4C and Table S4).

## Discussion

In our work, we found there was none of statistically difference of *EED* expression among LAML patients and normal subjects, but *EED* was significantly increased in DLBC patients compared with normal subjects. In LAML, expression of *EED* in LAML M3 subtype (also named as acute promyelocytic leukemia, APL) was obviously decreased compared with other LAML subtypes. In the TCGA clinical dataset, LAML patients were grouped into poor prognosis group, favorable prognosis group and intermediate/normal prognosis group according to the CALGB cytogenetics risk category. *EED* expression was significantly up-regulated in CALGB poor prognosis group compared with CALGB favorable prognosis group. Moreover, higher expression level of *EED* gene predicts shorter overall survival in LAML patients.

Our findings echoed some published articles. High expression of *EZH2* gene is involved in extramedullary infiltration in LAML and silencing of *EZH2* suppresses cell proliferation/migration and increases cell apoptosis in LAML (Zhu et al. 2016). Suppression of PRC2 subunits EED, SUZ12 or EZH1/EZH2 results in proliferation arrest and differentiation of leukemia cells (Shi et al. 2013). However, a set of articles involved in mice model demonstrated *EED* inactivation might be potential etiology of leukemia. Ikeda et al., report that *Eed* haploinsufficiency induces hematopoietic dysplasia and *Eed* heterozygous mice is susceptible to malignant transformation and developed leukemia in cooperation with *Evi1* overexpression (Ikeda et al. 2016). Ueda et al., demonstrate that *EED^I363M^* heterozygotes increases in the clonogenic capacity and bone marrow repopulating activity of hematopoietic stem/progenitor cells and is susceptible to leukemia through depressing *Lgals3*, encoding a multifunctional galactose-binding lectin (Ueda et al. 2016). Based on our work and published study, we guess *EED* might be a double-sword in LAML that loss-of-functions of *EED* contributes to LAML tumorigenesis and increased expression of *EED* predicts poor prognosis in LAML.

In order to investigate the regulatory network associated with *EED* expression, the *EED* co-expressed genes were identified and the KEGG pathway was enriched. In our work, we found, the co-expressed genes were significantly enriched in Fanconi anemia pathway, cell cycle, Oocyte meiosis, DNA replication, which also had the high connectivity with those co-expressed genes.

Fanconi anemia (FA) is a rare autosomal recessive cancer-prone inherited bone marrow failure syndrome, the major complications of which are aplastic anemia, acute myeloid leukemia (LAML) and myelodysplastic syndrome (MDS) (Alter 2014). *BRCA1* and *BRCA2* are DNA repair associated proteins. FA patients carry *FANCD1*/*BRCA2* mutations are susceptible to cancer risk, including leukemia, brain tumor or combinations (Alter 2014; Biswas et al. 2011). In our work, Fanconi anemia proteins, including *FANCB*, *FANCD2*, *FANCI*, *FANCL* and *FANCM*, co-expressed with *EED* and enriched in Fanconi anemia pathway. It is reported that hyper-methylation of *FANCL* promoter regions are found in sporadic acute leukemia (Hess et al. 2008). In addition, genetic variant of human homologous recombination-associated gene RMI, which co-expressed with *EED* and enriched in Fanconi anemia pathway, significantly increase the risk of LAML (Broberg et al. 2007).

Cell cycle is commonly dysregulated in various types of tumor. In our work, cell cycle proteins, *CCNB1*, *CCNB2*, *CCNE2*, *CDC27*, *CDC6*, *CDC7*, *CDK1*, co-expressed with *EED* and significantly enriched in cell cycle pathway. LAML patients with higher level of nuclear CDK1 in their leukemic blasts had the poorer clinical outcome and CDK1 modulate the level of p27 (kip) and AKT phosphorylation in response to ATRA treatment(Hedblom et al. 2013). The recent Phase I study of the novel CDK1 inhibitor terameprocol in patients with leukemia shows that 5 out 16 patients with advanced LAML or MDS have stable disease greater/equal to 2 months though reducing the expression of CDK1 and phosphor-AKT (Tibes et al. 2015).

*AURKA* and *PPP2R5E*, co-expressed with *EED*, were significantly enriched in oocyte meiosis pathway. High expression of *AURKA* and *AURKB* is associated with unfavorable cytogenetic abnormalities and high blood cell counts in LAML patients (Lucena-Araujo et al. 2011). Over-expression of *AEG-1* is essential for carcinogenesis role in human LAML through up-regulation of Akt mediated by *AURKA* activation (Long et al. 2013). *PPP2R5E* is commonly down-regulated and influence on the oncogenic potential of leukemia cells (Cristobal et al. 2013).

In our study, we found *EED* expression was significantly increased in DLBC compared with normal subjects and it was also significantly increased in DLBC compared with LAML subjects. PRC2 and H3K27me3 might paly opposite roles in LAML and DLBC. Decreased *EZH2* predicts poor prognosis in LAML and high abundance of H3K27me3 predicts poor prognosis in DLBC (Oh et al. 2014; Shi et al. 2013). Increased *EED* expression might paly essential roles in DLBC tumorigenesis. *EED* mRNA expression was negatively correlated with DNA methylation in DLBC, which indicated that high expression of *EED* might be due to hypo-methylation in DLBC. In actually, there are cross-talk between DNA methylation and histone methylation in genes (Cedar & Bergman 2009). Different epigenetic factors synergistically regulate activation and silencing of gene expression in various biological processes. Our study displayed that *EED* expression was not associated clinical stage and prognosis of DLBC, which was consistent with published article that PRC2 expression is not correlated with OS and RFS of DLBC and high abundance of H3K27me3 predicts shorter OS and RFS in DLBC patients (Oh et al. 2016; Oh et al. 2014).

*ARPC5*, *RELT* and *DPP3* (data not shown) had the most strongly positive correlation with *EED* expression and *NR1D2*, *PAIP2B* and *GPD1L* (data not shown) had the most strongly negative correlation with *EED* expression in DLBC. It is reported that *DPP3* is overexpressed in breast cancer and increased mRNA expression of predict unfavorable prognosis in breast cancer patients (especially in ER^+^ breast cancer) (Lu et al. 2017). *NR1D2* regulates glioblastoma cell proliferation and motility (Yu et al. 2018). The significance of abovementioned genes in DLBC tumorigenesis and prognosis need to be further investigated in our future work.

The genes co-expressed with *EED* in DLBC were significantly enriched three pathways, including Pathogenic Escherichia coli infection, Shigellosis and Legionellosis, involved in pathogen infection and immunity response, which echoes the pathogenesis of DLBC. In fact, infectious agents have been implicated as the cause of non-hodgkin lymphoma (NHL). There are three types of pathogens that cause NHL, the first type is viruses that can directly transform lymphocytes, such as EBV, HPV8, and HTLV-1, the second type is HIV virus, which that cause loss of CD4+T cells and indirectly induce NHL, the third type is viruses that induce NHL through chronic immune stimulation, such as HCV(Engels 2007). DLBC with EB infection is associated with the rapidly deteriorating clinical course with treatment response, survival and progression-free survival (Park et al. 2007).

In addition, *ZNF252P* and *XRCC5* were the shared genes among *EED* co-expressed genes in LAML and DLBC. *ZNF252P* was positively correlated with *EED* expression in LAML; however, it was negatively correlated with *EED* expression in DLBC. *XRCC5* encodes X-ray repair cross complementing 5 and functions as modulating chromosomal stability. It is down-regulated both in non-small cell lung cancer and breast cancer with hyper-methylation in promotor region; the SNPs in *XRCC5* promoter region might be susceptible to breast cancer (Lee et al. 2007; Rajaei et al. 2014).

*EED* gene possesses potential application value in the prevention and treatment of DLBC. In fact, A breakthrough has been made in DLBC targeted therapy by using an EED protein inhibitor. Compound 43, a small molecular compound, allosterically inactivates the methyltransferase activity of PRC2 complex through inhibiting with its adaptor protein of EED, which contribute to decrease tumor size in xenograft model of lymphoma (Yang & Wang 2017). In the preclinical study, compound 43 have the roubst anticancer efficacy based on oral administration (Huang et al. 2017). Therefore, targeted inhibition of the function of EED protein is an effective anticancer strategy.

## Acknowledgements

The authors thank Shanghai Jiao Tong University’s support.

---

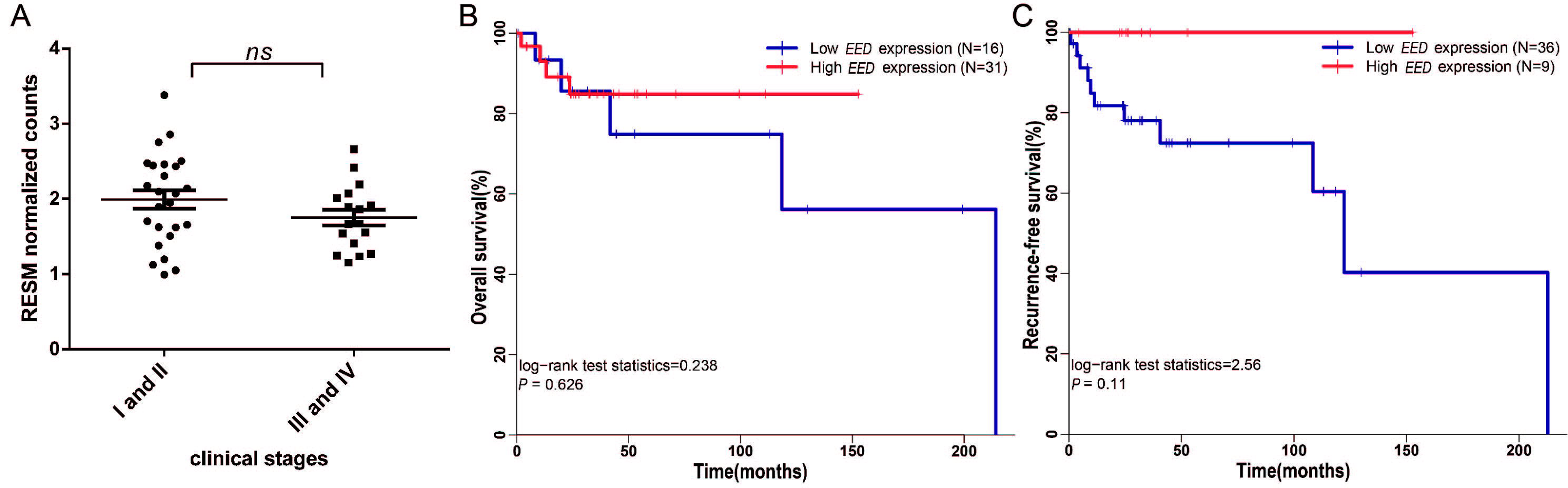

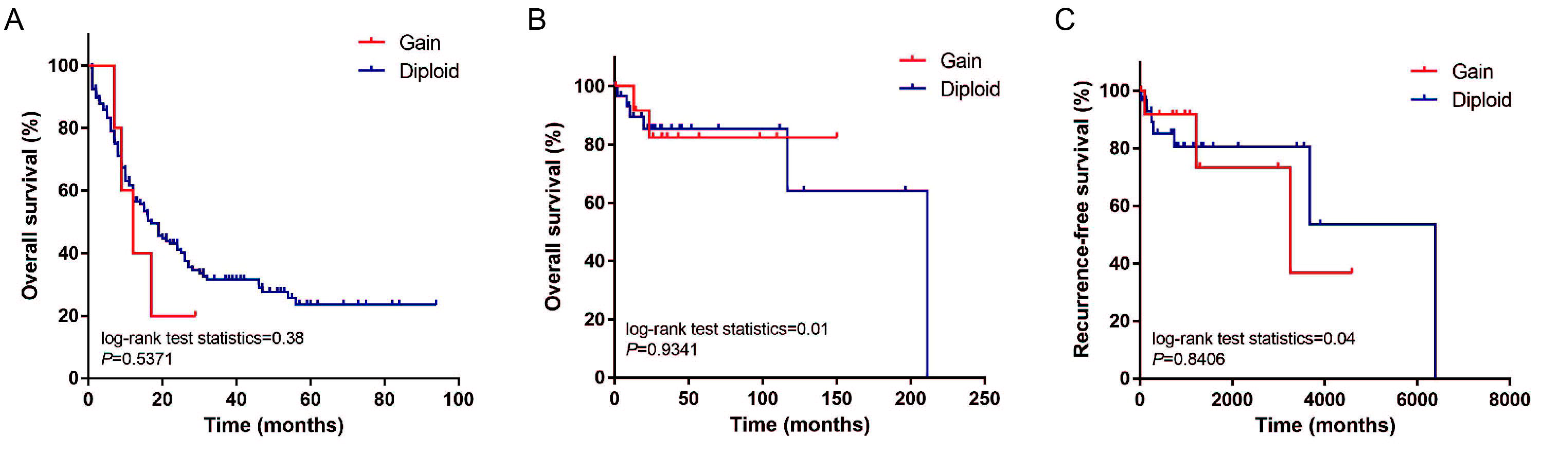

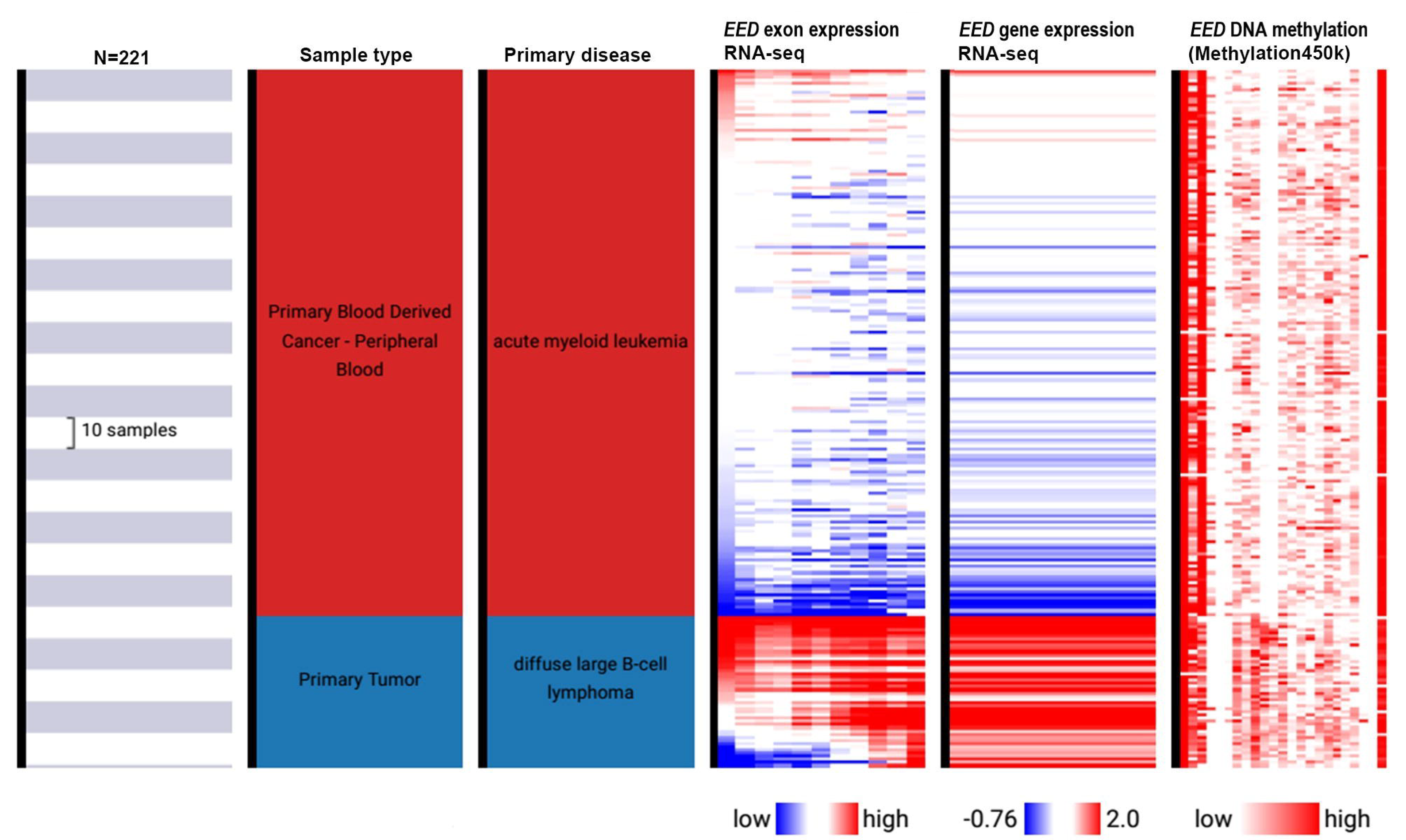

